# Adipocyte deletion of the RNA binding protein HuR induces cardiac hypertrophy and fibrosis

**DOI:** 10.1101/2021.01.19.425776

**Authors:** Adrienne R. Guarnieri, Sarah R. Anthony, Anamarie Gozdiff, Lisa C. Green, Sam Slone, Michelle L. Nieman, Perwez Alam, Joshua B. Benoit, Onur Kanisicak, Michael Tranter

## Abstract

Adipose tissue continues to gain appreciation for its broad role as an endocrine organ, and disruptions in adipose tissue homeostasis plays a central role in cardiovascular physiology. We have previously shown that expression of the RNA binding protein HuR in adipose tissue mediates energy expenditure, but the potential cardiovascular impacts of this finding have not been explored. We show here that adipose tissue-specific deletion of HuR (Adipo-HuR^-/-^) is sufficient to induce the spontaneous development of cardiac hypertrophy and fibrosis. Hearts from Adipo-HuR^-/-^ mice have increased left ventricular (LV) ejection fraction, rate of pressure generation, and LV posterior wall thickness that is accompanied by an increase in LV/body weight ratio and hypertrophic gene expression. Furthermore, Adipo-HuR^-/-^ hearts display increased fibrosis by picrosirius red staining and periostin expression. To identify underlying mechanisms, we applied both RNA-seq and weighted gene co-expression network analysis (WGCNA) to define HuR-dependent changes in gene expression as well as significant relationships between adipose tissue gene expression and LV mass. RNA-seq results demonstrate a significant increase in pro-inflammatory gene expression in the subcutaneous white adipose tissue (scWAT) from Adipo-HuR^-/-^ mice that is accompanied by an increase in serum levels of both TNF-α and IL-6. WGCNA identified a significant enrichment in inflammation, apoptosis/cell death, and vesicle-mediated transport genes among those whose expression most significantly associated with CVD in Adipo-HuR^-/-^. In conclusion, we demonstrate that the loss of HuR expression in adipose tissue promotes the development of cardiac hypertrophy and fibrosis, potentially through modulation of inflammation and vesicle-mediated transport in scWAT.

**NEW AND NOTEWORTHY:** This work demonstrates the spontaneous development of cardiac hypertrophy and fibrosis upon adipose tissue-specific deletion of the RNA binding protein HuR that appears to be mechanistically driven by HuR-dependent changes in inflammatory and extracellular vesicle transport mediating genes in the subcutaneous white adipose tissue. These results suggest that loss of HuR expression in adipose tissue in obesity, as demonstrated in mouse and humans by our group and others, may contribute to obesity-mediated CVD.

## INTRODUCTION

Obesity and metabolic syndrome are known co-morbidities of cardiovascular disease (CVD), and as the prevalence of obesity in the U.S. continues to climb, understanding of the causal relationship between disruption in adipose tissue homeostasis and CVD is of critical significance. Obesity and metabolic syndrome are both driven by imbalances in energy consumption, storage, and expenditure, and are marked by an increase in lipid storage and disruption of adipose tissue homeostasis. When energy intake exceeds expenditure, excess caloric energy is stored in white adipose tissue (WAT) as triglycerides. In contrast, beige and brown adipose tissue (BAT) drive energy expenditure through uncoupled thermogenic metabolism in their conserved evolutionary role in thermoregulation. Functional BAT, previously only believed to be in infants and children, was recently discovered in adult humans (12, 80) and is considered a potential therapeutic target in obesity due to its high metabolic potential, which is capable of accounting for up to 20% of total whole-body energy expenditure (65, 75, 76).

Adipose tissue has long been linked to CVD through its central role in obesity, but work over the past two decades has established a clear endocrine organ role for adipose tissue that is dependent on both adipocyte composition (e.g. WAT vs. BAT) as well as anatomical location (e.g. visceral vs. subcutaneous) (2). Specifically, accumulation of WAT tends to be more pro-inflammatory in nature, especially under conditions of chronic energy surplus, and exerts deleterious effects on cardiovascular health (19, 55, 59). On the other hand, the presence of metabolically active BAT has been demonstrated to positively correlate with myocardial health (2, 63, 72, 74).

Human antigen R (HuR) is a widely-expressed RNA binding protein that interacts with specific AU-rich domains in target mRNAs and directly regulates expression by modulating mRNA stability and/or translation (14). Our recently published work was among the first to show the *in vivo* role of HuR on adipocyte physiology (1). Two other groups have also recently shown roles for HuR in adipogenesis and lipolysis (48, 69). We specifically showed that adipocyte-specific deletion of HuR in mice (Adipo-HuR^-/-^ mice) resulted in a reduction in both WAT and BAT mass that was accompanied by a profound deficit in adaptive thermogenesis when subjected to acute cold challenge (1). Interestingly, Li et al recently showed an inverse correlation between HuR expression in subcutaneous WAT (scWAT) and obesity in human patients (48), suggesting a clinical link for HuR expression in metabolic syndrome and raising the question as to whether loss of HuR expression in adipose tissue may play a functional role in obesity associated co-morbidities such as CVD. Prior to showing a role for HuR in adipocyte function, we independently demonstrated a role for HuR expression in cardiomyocytes during the progression of pathological cardiac hypertophy (24, 70). However, in light of the growing interest in the adipose tissue-to-myocardium signaling axis, potential HuR-mediated cross-talk between adipose tissue and the myocardium has not yet been explored.

Here, we demonstrate that adipocyte-specific deletion of HuR mediates the spontaneous development of left ventricular cardiac hypertrophy and fibrosis in mice that appears to be mechanistically driven by an increase in inflammatory gene expression in subcutaneous white adipose tissue.

## MATERIALS AND METHODS

### Animal Models

All mouse studies were approved by the University of Cincinnati Institutional Animal Care and Use Committee (IACUC). HuR floxed mice were described by Ghosh et al (22) and obtained from Jackson Labs (stock # 021431). To generate an adipose-specific HuR deletion model (*Adipo-HuR*^*-/-*^), HuR-floxed mice were crossed with mice harboring an adiponectin-specific Cre recombinase transgene (*AdipoQ-Cre*) (Jackson Labs, stock # 010803). Non-floxed *Adipo-cre*^*+*^ littermates were used as controls. Mice were housed in the University of Cincinnati vivarium at 23°C. All data presented is from 18-22 week old male mice.

### Echocardiography

All echocardiographic studies were performed as previously described (24). Briefly, mice were sedated with isoflurane and body temperature was maintained at 37°C during imaging. Using a Vevo 2100, parasternal images were obtained in long axis in two-dimensional mode and motion (M)-mode for quantification. These were then analyzed using VevoStrain software (Vevo 2100, v1.1.1 B1455, Visualsonic, Toronto, Canada).

### LV Pressure Catheterization

As a terminal procedure, mice were weighed and anesthetized using inhaled isoflurane. Terminal development of LV pressure was determined using a 1.2-French catheter pressure transducer (Scisense FTH-1211B-0018) advanced through the carotid into the LV as previously described (17, 51).

### Tissue and plasma processing

At time of euthanasia, mice were sedated with isoflurane and blood collected into 3.2% sodium citrate to collect plasma via centrifugation at 4,000 x g at room temperature, which was snap frozen. Mice were subsequently euthanized via thoracotomy and tissues were removed, weighed, and flash frozen in liquid nitrogen for further analysis. Plasma catecholamine (epinephrine, norepinephrine; Abnova KA1877), IL-6 (Bioss BSKM1004) and TNF-α (Bioss BSKM1002) levels were assessed using ELISA kits according to the manufacturers’ instructions.

### Histological Analysis

Frozen hearts were embedded in OCT, sectioned, and stained using Picrosirius red (Polysciences, Inc) as previously described (29). Briefly, slides were air dried, immersed in xylene for 10 minutes, and hydrated through decreasing concentrations of ethanol (100%, 96%, 80% and 70%, 10 seconds each). They were then stained according to the manufacturer’s instructions, dehydrated through increasing concentrations of ethanol, cleared in xylene, and mounted with Permount under a glass coverslip.

### RNA Isolation and qPCR

RNA was isolated using a Macherey-Nagel NucleoSpin RNA kit and cDNA was synthesized using iScript RT Supermix (Bio-Rad). Samples were run on a Bio-Rad CFX96 Touch to assess mRNA levels of BNP, RCAN, and 18S. Primers are as listed: BNP, F: 5’-AAGTCCTAGCCAGTCTCCAGA-3’, R: 5’-GAGCTGTCTCTGGGCCATTTC-3’; RCAN, F: 5′-GGGCCAAATTTGAATCCCTCTTC-3′, R: 5′-GGAGCCAGGTGTGAACTTCC-3′; 18S, F: 5’-AGTCCCTGCCCTTTGTACACA-3’, R: 5’-CCGAGGGCCTCACTAAACC-3’.

### Protein Isolation and Western blotting

Total protein was isolated from crushed tissue in RIPA buffer with 0.5 mM DTT, 0.2 mM sodium-orthovanadate, and a protease inhibitor mixture tablet (Complete mini; Roche Applied Science). 25 ug of protein extract per lane was separated on a 10% polyacrylamide gel and transferred to a nitrocellulose membrane. Blocking was performed for 1 h at room temperature using 5% BSA milk in 0.1% Tween 20, tris-buffered saline (T-TBS). Primary antibodies for Periostin (Novus Biologicals NBP1-30042), PERK (Cell Signaling, 3192), and GAPDH (Santa Cruz Biotechnology sc25778) were incubated overnight at 4°C, and secondary antibodies were incubated for 1–2 h at room temperature in T-TBS. Images were captured and analyzed using a ChemiDoc Imaging System and ImageLab software (BioRad).

### RNA Sequencing and Weighted Gene Co-Expression Network Analysis (WGCNA)

RNA was isolated from cardiac and adipose tissue as previously described (1, 70) and RNA sequencing was done by the Cincinnati Children’s Hospital Medical Center (CCHMC) DNA Sequencing and Genotyping Core. Sequence read mapping, principal component analysis, and differential expression analysis was done using CLC Genomics Workbench (v. 20.0.2, Qiagen) as previously described (1, 24). Gene ontology analysis was done using the NIH DAVID Bioinformatics Functional Annotation Tool with an EASE score/P-value threshold of 0.05 as previously described (35, 36, 77). Significant differential expression between groups was defined as an absolute fold change of greater than or equal to 1.5 and a false discovery rate adjusted (FDR) p-value of less than or equal to 0.05. For WGCNA, co-expression network construction, module detection, association of expression modules with phenotype data, and identification of node genes withing expression modules was done as previously described (46, 89). Node genes were defined as the top 10 genes with the highest sum of absolute module membership and gene significance values and a gene significance value of > 0.85. All RNA-seq data is available in the NCBI Gene Expression Omnibus (https://www.ncbi.nlm.nih.gov/geo/) (GDSXXX).

## RESULTS

### Cardiac function in Adipo-HuR^-/-^ mice

Examination of cardiac function in 20-week old mice by echocardiography showed an increase in left ventricular (LV) ejection fraction (Fig. 1A), decreased LV volume at systole (along with a strong trend to decrease in diastole; Figs. 1B-C), and increased LV posterior wall thickness (Fig. 1D) in hearts from Adipo-HuR^-/-^ mice compared to littermate controls. Invasive hemodynamic assessment via LV pressure catheterization also shows an increased rate of positive and negative pressure generation in Adipo-HuR^-/-^ hearts (Fig. 1E-F). As we have previously shown a decrease in body weight in Adipo-HuR^-/-^ mice (1), cardiac structural changes have been normalized to body weight. Together, these data are indicative of a state of compensated cardiac hypertrophy in Adipo-HuR^-/-^ mice.

**Figure 1.**
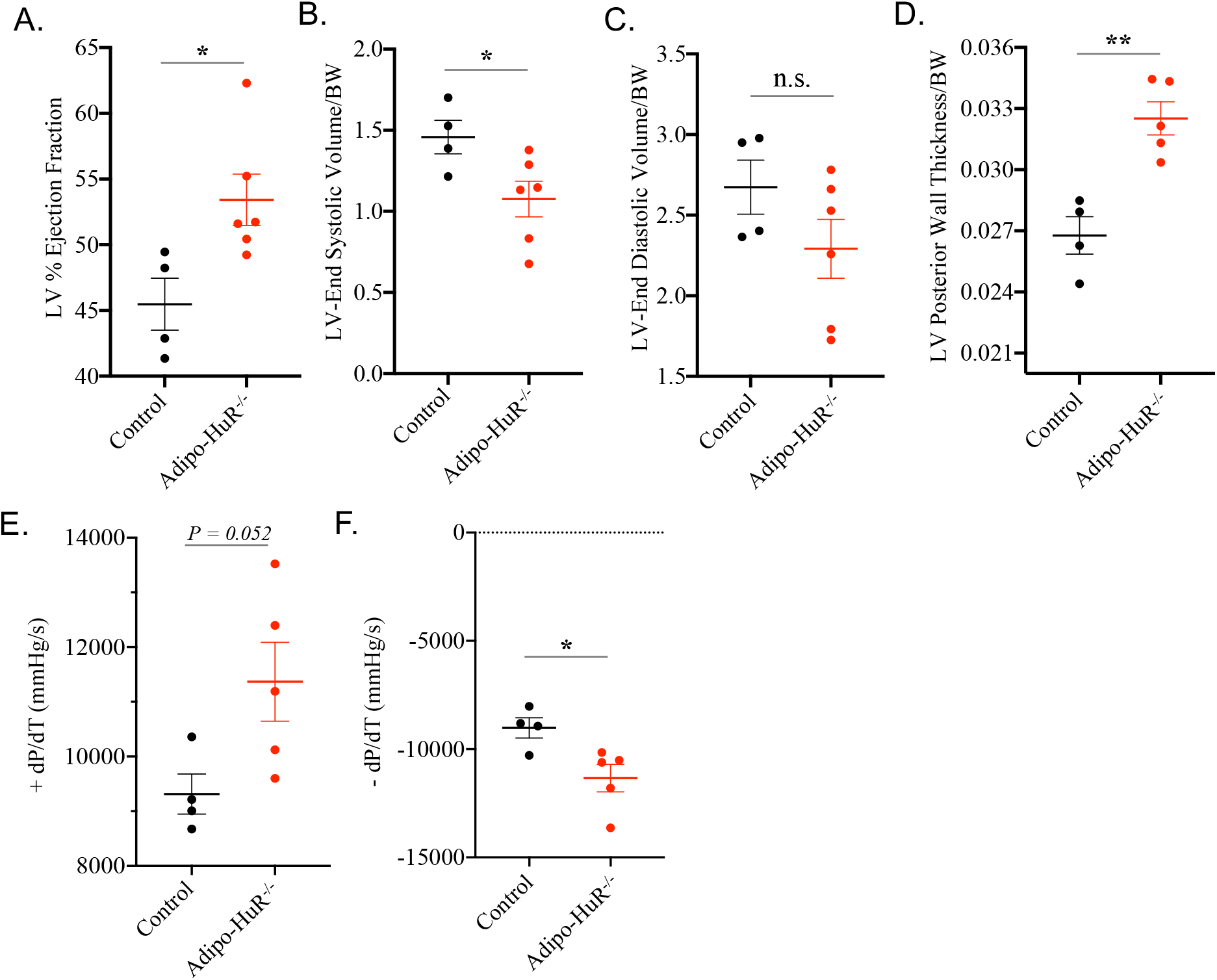
Functional assessment of cardiac function in hearts from mice with an adipocyte-specific deletion of HuR. Echocardiographic analysis shows that hearts from Adipo-HuR^-/-^ mice have increased %EF (A), with decreased LV chamber volume (B-C), and increased LV posterior wall thickness (D). LV pressure catheterization shows an exacerbation in the rates of positive and negative developed pressure (E-F). n ≥ 4 per group. *P ≤ 0.05; ** P ≤ 0.01.

### Molecular assessment of cardiac hypertrophy and fibrosis

First, LV mass relative to body mass was found to be significantly increased in Adipo-HuR^-/-^ mice (Fig. 2A), and qPCR showed increased expression of the hypertrophic marker genes BNP and RCAN (Fig. 2B-C). Next, assessment of cardiac fibrosis via picrosirius red collagen staining found increased interstitial fibrosis throughout the myocardium of Adipo-HuR^-/-^ mice (Fig. 2D-E). This was accompanied by a significant increase in protein expression of periostin, a marker for active myofibroblast ECM remodeling, and the ER-stress marker PERK (Fig. 2F-H). Thus, molecular signaling within the myocardium was found to be consistent with pathological hypertrophy.

**Figure 2.**
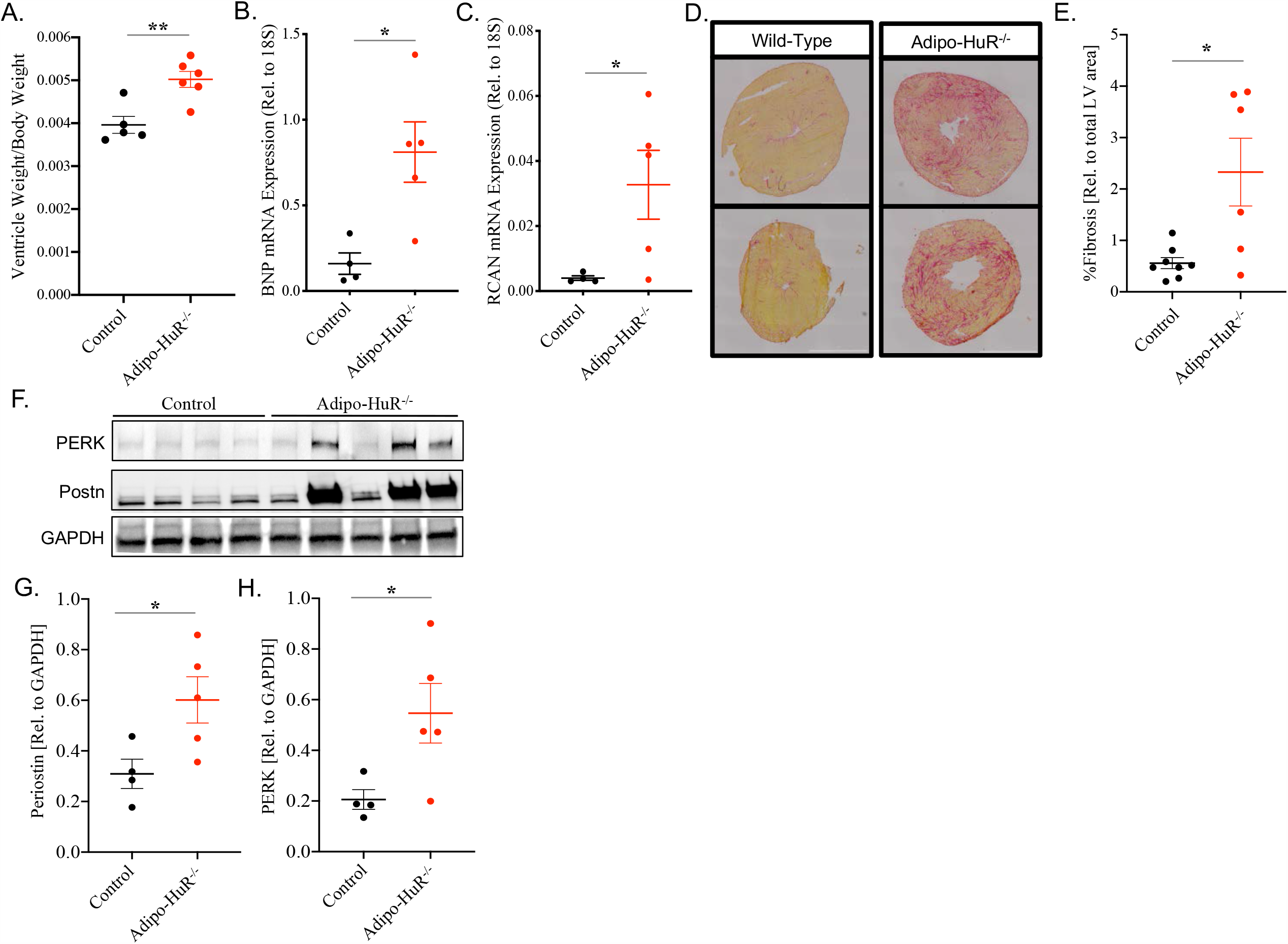
Molecular assessment of cardiac hypertrophy and fibrosis. Adipo-HuR^-/-^ have increased ventricle weight to body weight ratios compared to control mice (A). Gene expression of the hypertrophic markers BNP and RCAN were also increased in Adipo-HuR^-/-^ mice (B, C). Hearts from Adipo-HuR^-/-^ mice also display increased fibrosis compared to controls as shown by representative picrosirius red staining (D, E). Western blotting shows a significant increase in PERK and periostin protein expression in Adipo-HuR^-/-^ hearts (F-H). N > 4 per group. *P ≤ 0.05. **P ≤ 0.01.

### Circulating catecholamine levels

Adrenergic signaling response to cold activates thermogenic pathways, but also mediates increased cardiac contractility upon acute exposure and pathological cardiac hypertrophy with prolonged exposure (18). Thus, to rule out adrenergic-induced cardiac pathology as a result of cold-intolerance in Adipo-HuR^-/-^ mice, we show that plasma levels of epinephrine and norepinephrine are not increased (Fig. 3A and B, respectively). This demonstrates that the cardiac hypertrophy and fibrosis observed in Adipo-HuR^-/-^ mice is not driven by increased adrenergic signaling.

**Figure 3.**
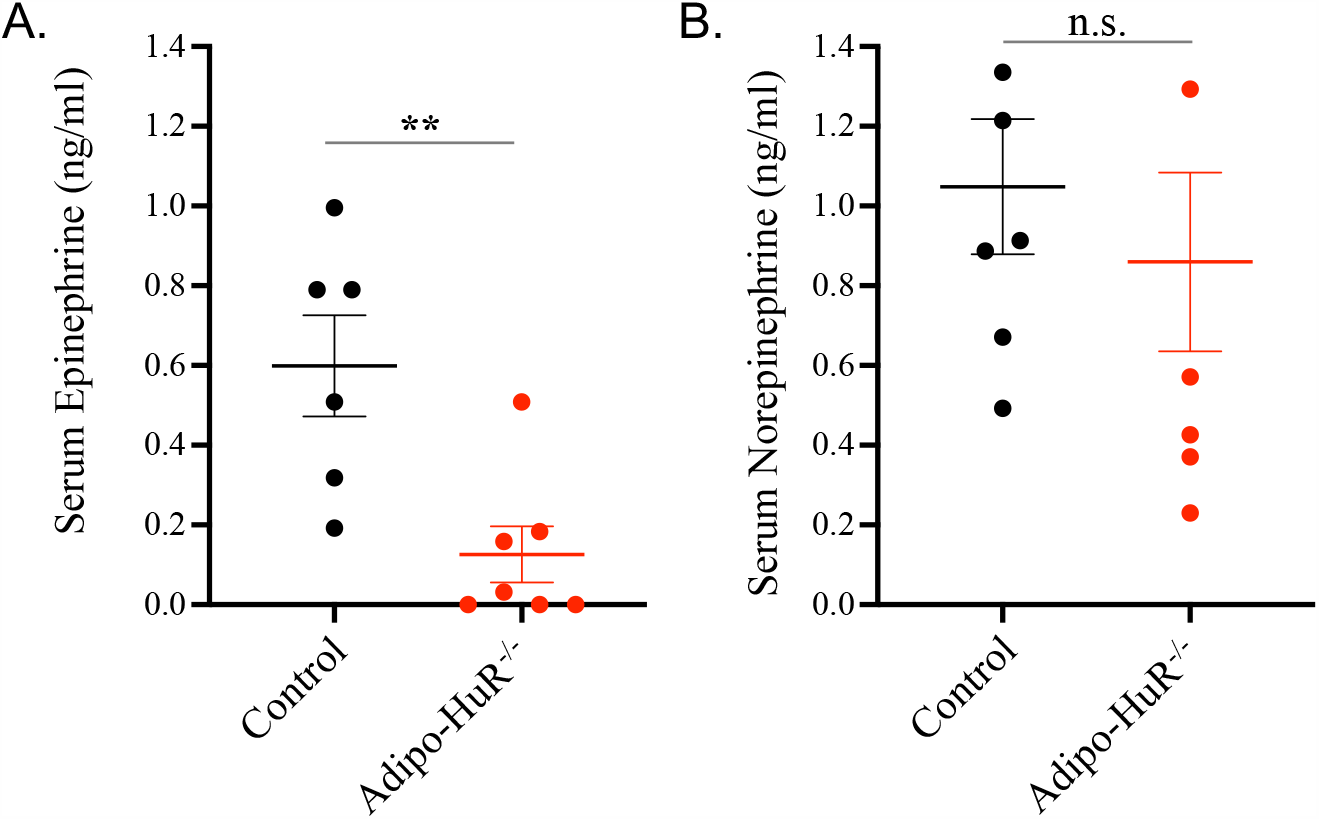
Serum levels of epinephrine (A) and norepinephrine (B) as determined by ELISA. N ≥ 5 per group. **P ≤ 0.01.

### RNA-seq assessment of cardiac gene expression

Transcriptome-wide RNA-sequencing was used to identify the full breadth of cardiac gene expression changes upon HuR deletion in adipose tissue. Principal component analysis showed a clear distinction between cardiac transcriptomes in Adipo-HuR^-/-^ mice and littermate controls (Fig. 4A). We identified a total of 476 significantly dysregulated genes, as defined by a fold change of ≥ 1.5 and a false discover rate (FDR)-corrected *P* value of ≤ 0.05. Of these significant genes, 218 were significantly down-regulated and 258 were significantly up-regulated relative to expression in control hearts. Gene ontology (GO) analysis identified a significant enrichment in GO terms for inflammatory gene expression among significantly upregulated genes in hearts from Adipo-HuR^-/-^ mice (Fig. 4B). In addition to an increase in many individual pro-inflammatory genes, RNA-seq analysis also confirmed a significant increase in pro-fibrotic and hypertrophic genes (Fig. 4C).

**Figure 4.**
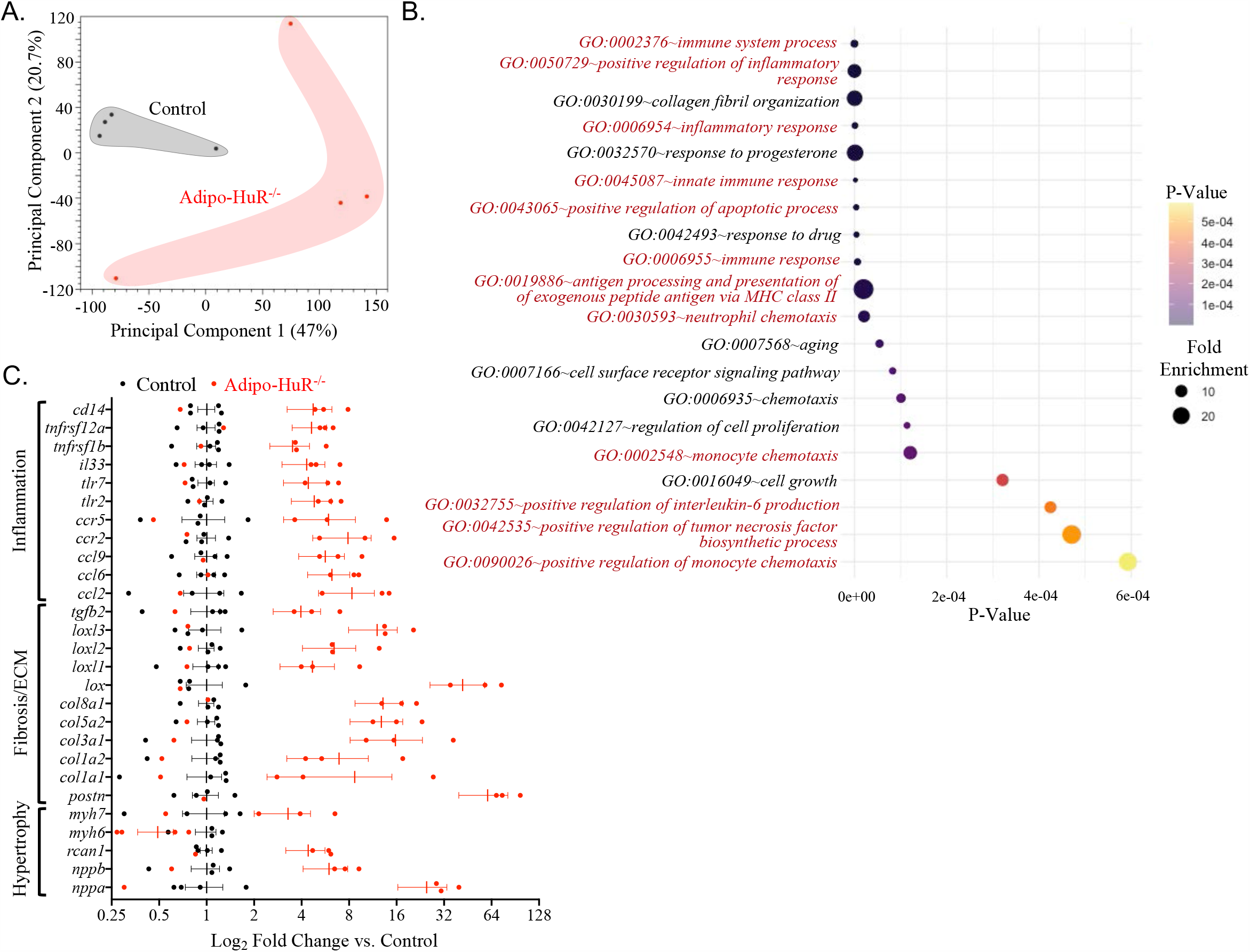
RNA-Seq analysis of gene expression in control and Adipo-HuR^-/-^ hearts. Principal component analysis (PCA) plot shows clustering of gene expression patterns from control and Adipo-HuR^-/-^ hearts (A). Gene ontology (GO) analysis shows a significant enrichment in inflammation-associated genes, indicated in red (B). RNA-seq analysis shows a significant increase in hypertrophic, fibrotic, and inflammatory gene expression (C); all genes shown in panel C are significant as defined by fold change ≥ 1.5 and FDR P ≤ 0.05. N = 4 per group.

### RNA-seq assessment of adipose tissue gene expression

We previously reported RNA-seq analysis of HuR-dependent changes in BAT (1), but found nothing that would be obviously indicative of cardiovascular disease. Here, we again applied RNA-seq (with a statistical cutoff of a fold change of ≥ 1.5 and an FDR *P* value of ≤ 0.05) to identify transcriptome-wide HuR-dependent gene expression in scWAT. Results show that while HuR mediates expression of 588 genes in BAT, we found 1117 HuR-dependent genes in scWAT (Fig. 5A). Interestingly, many of the HuR-dependent gene expression changes in BAT were decreases in expression, but most HuR-dependent changes in scWAT were increases in expression, suggesting differing roles for HuR-mediated gene expression in the two tissues (Fig. 5B).

**Figure 5.**
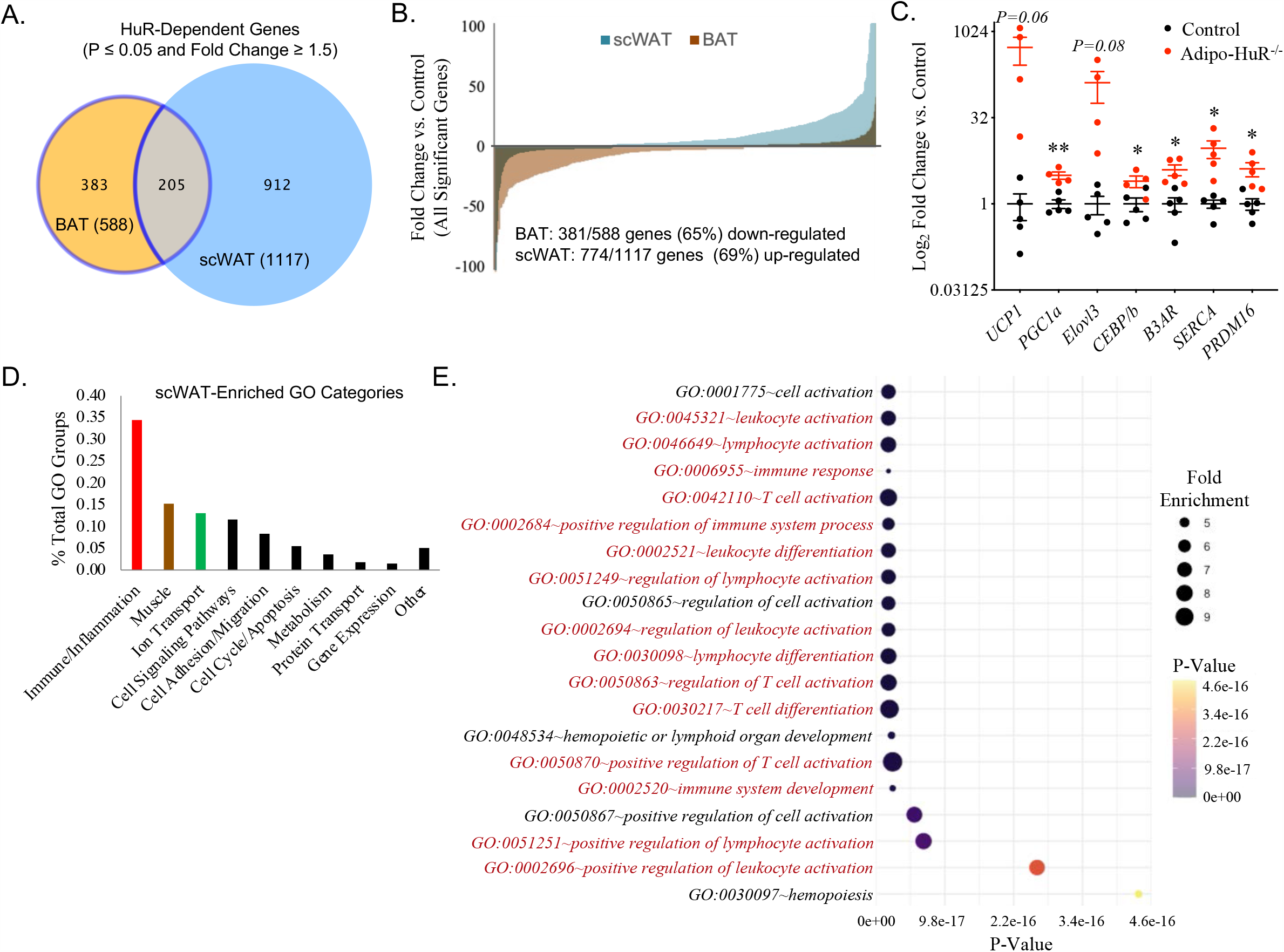
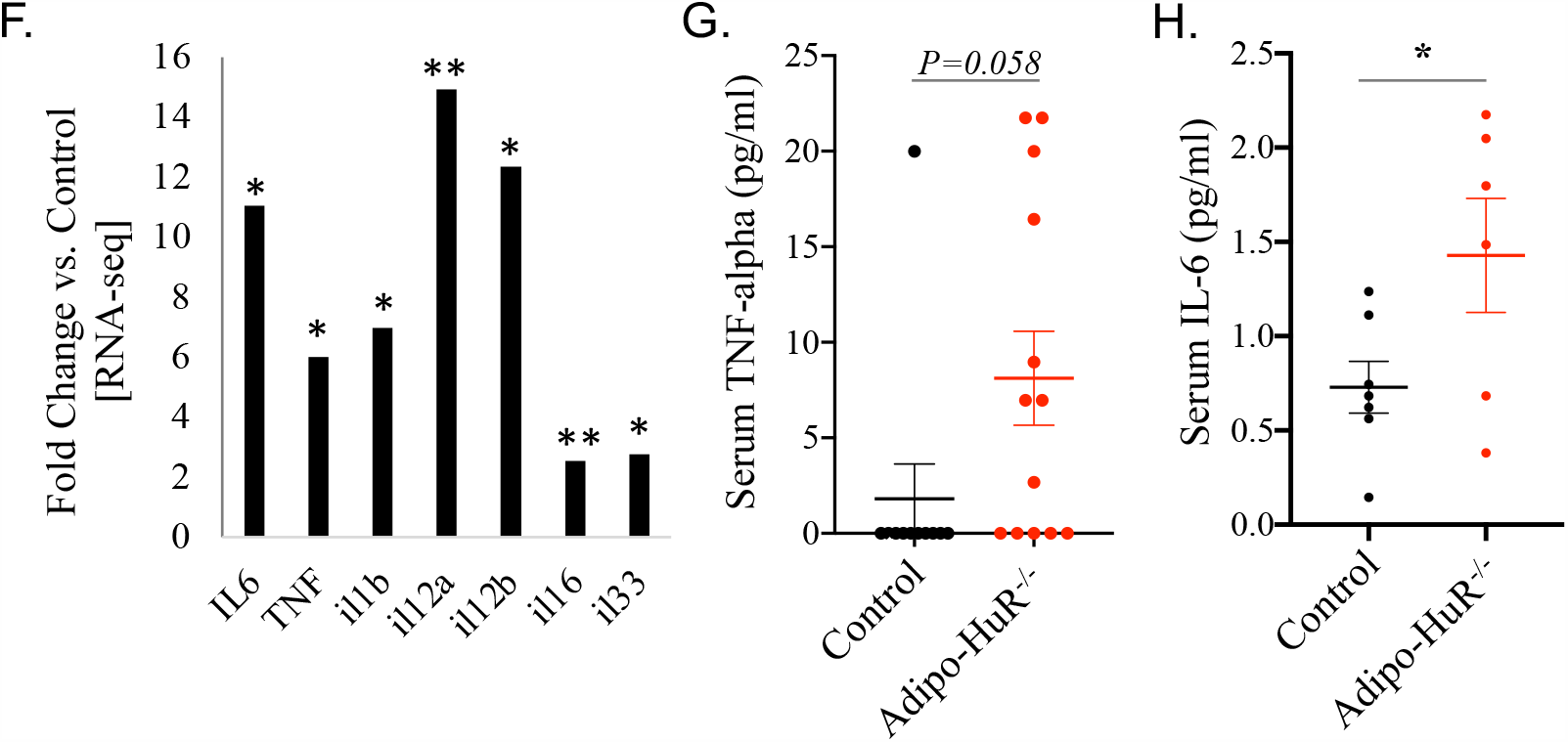
RNA-Seq analysis of gene expression in control and Adipo-HuR^-/-^ scWAT. Venn diagram of HuR-dependent changes in gene expression changes in BAT and scWAT shows a unique set of HuR-dependent genes in the two adipose depots (A). The majority (381/588; 65%) of the significant HuR-dependent changes in gene expression in BAT are down-regulated, while the majority (774/1117; 69%) are up-regulated in scWAT (B). Expression of beige adipocyte marker genes in scWAT (C). Gene ontology analysis showing classification by percent of all enriched HuR-dependent GO terms in scWAT (D) and the top 20 enriched HuR-dependent GO terms among genes with significantly increased expression (E). RNA-seq determined expression change of proinflammatory cytokines in scWAT that have previously been suggested to mediate cardiac pathology through secreted/paracrine signaling (F). N = 4 per group. Serum levels of TNF-α (G) and IL-6 (H) as determined by ELISA. N ≥ 5 per group. *P ≤ 0.05. **P ≤ 0.01.

Among the significant gene expression changes in scWAT, we observe an increase in many pro-thermogenic genes/beige adipocyte markers (Fig. 5C), which is not surprising given thermogenic deficiency previously reported in Adipo-HuR^-/-^ mice (1). Gene ontology (GO) analysis of HuR-dependent gene expression changes in scWAT show an overwhelming enrichment in pro-inflammatory mediators, as ∼33% of all significant GO terms are associated with inflammation/immune processes (Fig. 5D). Many of the top 20 enriched GO terms among HuR-dependent up-regulated genes in scWAT are inflammation associated GO terms (Fig. 5E). Among these increased inflammatory gene products are several pro-inflammatory cytokines which play a detrimental role in cardiac physiology (Fig. 5F) (13, 21, 26, 73). Importantly, we also show an increase in plasma levels of TNF-αand IL-6 (Fig. 5G and 5H, respectively), suggesting that this large increase in inflammatory gene expression in scWAT results in an increase in systemic inflammation.

### Association of adipose tissue expression networks with cardiac phenotype

To more specifically determine which gene expression changes within the scWAT are likely to underlie specific pathological changes in the heart, we applied weighted gene co-expression network analysis (WGCNA) (46, 89). Here, we applied WGCNA analysis to cluster the scWAT transcriptome into co-expressed gene modules, then associate each module with cardiac phenotype through a parallel analysis of module-phenotype correlation and average module eigengene significance within each module (Fig. 6A). After clustering the scWAT transcriptome into co-expressed gene modules independent of genotype (Fig. 6B), cardiac hypertrophy (left ventricle weight to body weight ratio; LVW/BW) was used as the specific cardiac phenotype of interest to identify scWAT gene expression associations. We identified three significant co-expression modules (*lightsteelblue1, magenta*, and *yellowgreen*) that passed criteria for both module-phenotype correlation (*P* ≤ 0.05; Fig. 6C) and average gene significance (0.75) relative to LV/BW (Fig. 6D). Overall significance of these modules was confirmed through linear correlation demonstrating a strong relationship between individual gene significance to phenotype and module membership (Fig. 6E).

**Figure 6.**
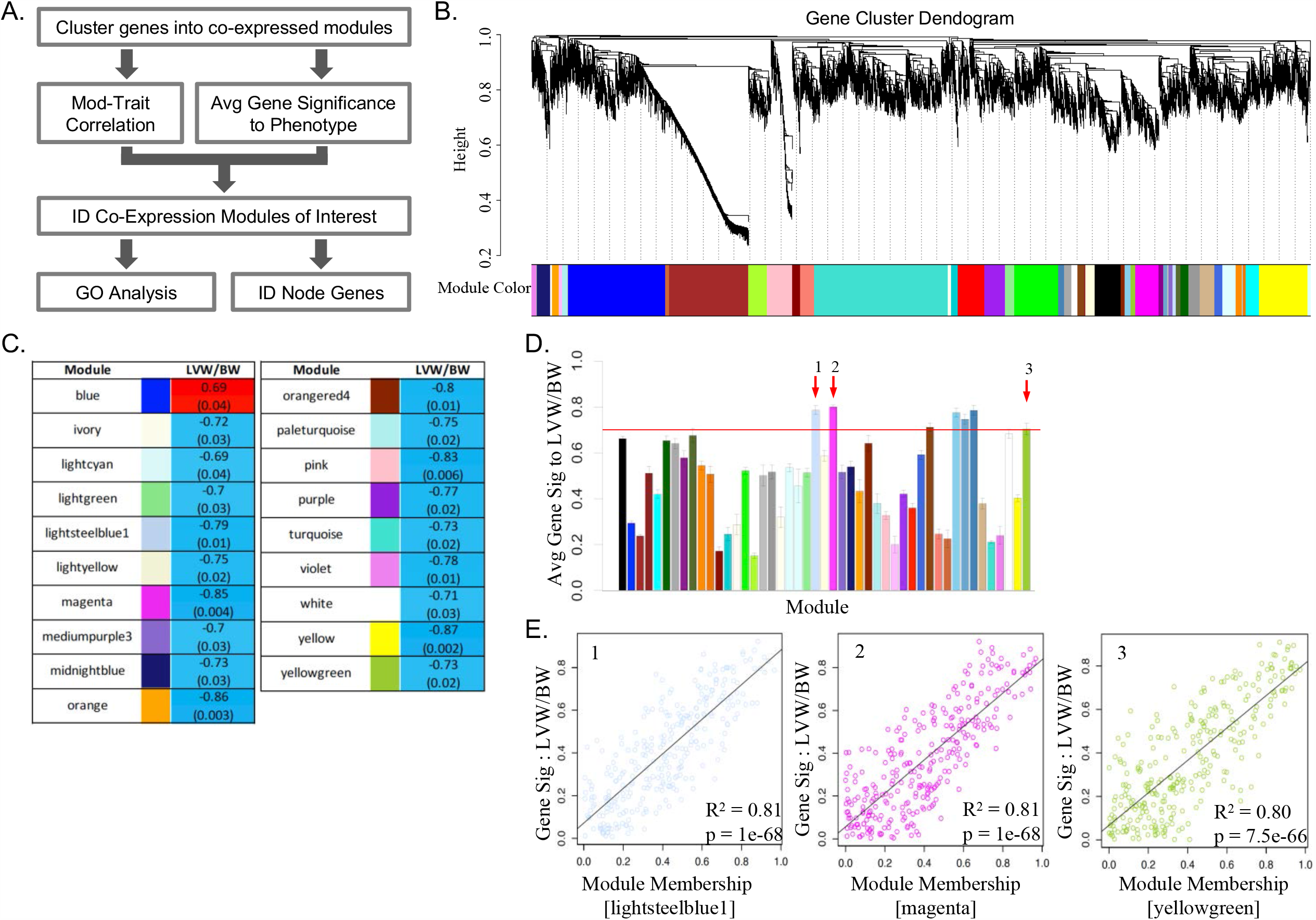
WGCNA association of scWAT gene expression with cardiac hypertrophy phenotype. Workflow used to identify and associate co-expressed gene modules with phenotype independent of genotype (A). Gene cluster dendogram of co-expressed gene modules (B). Correlation of each gene expression module with LVW/BW was used as a first filter to identify significant modules (C). Average gene signficance relative to LVW/BW was calculated for each module as a second filter for identification of significant modules (D). (E) Confirmation of a strong significant correlation between gene significance to LVW/BW and module membership for individual genes within the three identified modules that pass both correlation (C) and average gene significance (D).

Subsequent Gene ontology analysis of individual genes within these three co-expression modules revealed a significant enrichment for genes associated with apoptosis/cell death and vesicle mediated transport (Figure 7). Within each module, individual node genes, or those that have the highest gene significance and module membership score within each module, can be identified as those genes that have the highest likelihood of having a causal relationship with the phenotype. Congruent with the GO results, the majority of these node genes play a significant role in either inflammation, apoptosis, or vesicular mediated transport. A full list of these node genes, their individual association with LVW/BW, and relevant biological functions are found in Table 1. The association of apoptosis-related co-expression modules in scWAT with cardiac pathology is consistent with the inflammatory phenotype observed in both tissues, but these results also suggest paracrine signaling via extracellular vesicles as a potential mediator of scWAT-derived mediator of pathological cardiac hypertrophy.

**Table 1.** Node gene identification significant LVW/BW associated modules.

|  |  |  |  |  |
| --- | --- | --- | --- | --- |
| lightsteelblue1 | Symbol | Gene Sig LVW/BW | P-value | Relevant Biological Function |
|  | *mat2b* | 0.9242 | 0.0004 | Adipogenesis (1); Proliferation/Anti-apoptosis (2) |
|  | *sumf2* | 0.8968 | 0.0011 | IL-13 expression and secretion (3, 4) |
|  | *cuedc2* | 0.8895 | 0.0013 | Inflammatory signaling (5, 6, 7); myocardial I/R injury and oxidative stress (8) |
|  | *wdfy1* | 0.8819 | 0.0017 | TLR4 signaling (9); Endosome trafficking (10) |
|  | *mrpl27* | 0.8708 | 0.0022 | Experimental correlation with CVD in metabolic syndrome (11) |
|  | *pef1* | 0.8693 | 0.0023 | Ca-dependent ER-Golgi transport and collagen export (12) |
|  | *edf1* | 0.8620 | 0.0028 | Lipid metabolism (13); Myocyte hypertrophy (14); CaM/NFAT signaling during adipogenesis (15); PPARgamma transcriptional cofactor (16) |
|  | *tmem160* | 0.8577 | 0.0031 | Unknown; Clinical association with obesity risk (17, 18) |
|  | *edem1* | 0.8539 | 0.0034 | ATF6-mediated cell survival ER stress response (19) |
|  | *ifi27* | 0.8503 | 0.0037 | IFN-gamma mediated apoptosis (20-22) |
| magenta | Gene | Gene Sig LVW/BW | P-value | Relevant Biological Function |
|  | *Atp6v1c2* | 0.95 | 0.0001 | Acidification of intracellular compartments |
|  | *Pigs* | 0.95 | 0.0001 | Biosynthesis of glycolipid cell surface anchor |
|  | *Eno1* | 0.94 | 0.0002 | Identified in extracellular vesicles/exosomes (23-27); Promotes ECM degradation (28) and inflammation (29) |
|  | *Gltp* | 0.93 | 0.0003 | Vesicular trafficking, sphingolipid homeostasis, necroptosis (30) |
|  | *Rap1gap2* | 0.92 | 0.0004 | RAP1 GTPase signaling (31); Dense granule secretion in platelets (32) |
|  | *Tmem254c* | 0.92 | 0.0004 | Unknown |
|  | *Vldlr* | 0.91 | 0.0007 | Receptor-mediated endocytosis |
|  | *Commd10* | 0.91 | 0.0005 | NF-kappaB signaling (33, 34); Endocytosis (35); Ly6C(hi) monocyte inflammasome activity (36); Clinical GWAS correlation with inflammatory biomarkers (37) |
|  | *Fkbp1a* | 0.89 | 0.0012 | TGF-beta signaling (38); Ryanadine receptor-mediated calcium transport (39); Left ventricular non-compaction (40); Immunosuppressant in complex with rapamycin (41) |
|  | *Nln* | 0.88 | 0.0015 | Glucose tolerance/insulin sensitivity (42); Adipose expression associated with angiotensin-mediated weight gain (43) |
| yellowgreen | Gene | Gene Sig LVW/BW | P-value | Relevant Biological Function |
|  | *Bicdl1* | 0.91 | 0.0006 | Regulates secretory vesicle transport (44) |
|  | *Tmem62* | 0.91 | 0.0007 | Unknown |
|  | *Tmem97* | 0.89 | 0.0014 | LDLR endocytosis (45); Cellular cholesterol homeostasis (46) |
|  | *Mvb12b* | 0.87 | 0.0022 | Mediates sorting of ubiquitinated proteins to endosomal vesicles (47) |
|  | *Oas1c* | 0.86 | 0.0029 | SNP associated with diabetes susceptibility (48) |
|  | *Ccs* | 0.86 | 0.0030 | Copper homeostasis and oxidative stress (49) |

**Table 2.** Node gene identification within significant LVW/BW associated modules.

**Figure 7.**
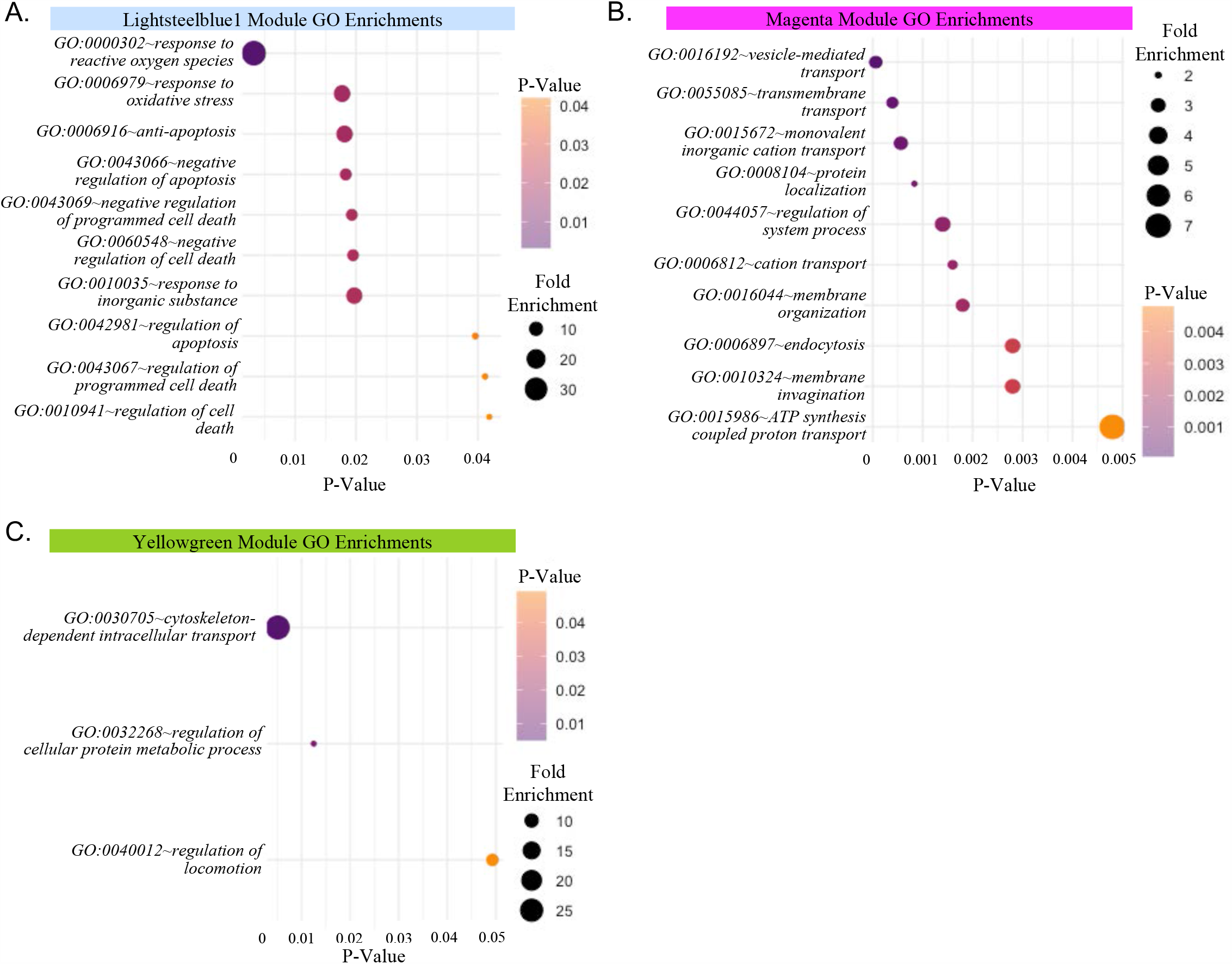
Enriched GO terms among genes from significant LVW/BW-associated co-expression modules. The lightsteelblue1 module (A) had exactly 10 significantly enriched GO terms. GO enrichments for the magenta module (B) represent the 10 most significantly enriched of 59 total signficant terms. The yellowgreen module (C) had only 3 significantly enriched GO terms.

## DISCUSSION

In this work, we demonstrate that hearts from Adipo-HuR^-/-^ mice develop cardiac hypertrophy with increased LV/BW ratio, LV wall thickness, and increase in hypertrophic marker genes. These mice also show an increase in cardiac fibrosis with increased expression of fibrotic (periostin) and ER stress (PERK) proteins. Increased PERK expression indicates activation of the unfolded protein response (UPR) and ER stress, which has been shown to be induced in multiple forms of CVD and is increased in failing human hearts (37, 90). Consistent with these results, RNA-seq analysis also shows a broad increase in hypertrophic, fibrotic, and inflammatory marker genes in these hearts confirming pathological hypertrophy and fibrosis. Perhaps most significant is that the cardiovascular pathology described in Adipo-HuR^-/-^ mice is driven via adipocyte-specific signaling in the absence of glucose intolerance, changes in circulating lipid levels, or obesity (all data presented are from lean mice maintained on control chow diet). We have previously shown that Adipo-HuR^-/-^ mice have no differences in glucose tolerance or plasma phospholipids, cholesterol, triglycerides, or non-esterified fatty acids (1).

Both scWAT inflammation and BAT dysfunction have been previously associated with CVD through disparate independent mechanisms (2). In our prior work, we demonstrated the role of adipocyte-specific HuR deletion on the function of BAT, including a full RNA-seq analysis of HuR-dependent gene expression in BAT (1), but found nothing that would suggest a CVD-mediating effect emanating from direct HuR-dependent changes in BAT. However, data presented here shows a large degree of gene expression in scWAT that appears to be the result of direct compensation for the loss of BAT-mediated thermogenesis. Indeed, many gene markers of thermogenesis suggesting the beiging of scWAT, such as *ucp1, elovl3*, and *cidea*, are significantly increased in scWAT from Adipo-HuR^-/-^ mice. Interestingly though, the majority of gene expression changes in scWAT, as categorized by gene ontology analysis, were associated with immune and inflammatory processes. This increased inflammatory gene expression in scWAT is accompanied by a systemic increase in circulating IL-6 and TNF-α, both of which have been shown to mediate CVD (26, 56, 92).

To expand upon the standard RNA-seq analysis and identify potentially unique gene expression changes within the scWAT that are likely to underlie specific pathological changes in the heart, we applied weighted gene co-expression network analysis (WGCNA) (46, 89). WGCNA has been shown to be a powerful tool to directly correlate gene expression changes with phenotype and is an excellent predictor of causal relationships (28, 81, 83). Gene ontology analysis of individual genes within co-expressed gene modules showing a positive association with cardiac hypertrophy revealed two central themes: apoptosis/cell death and vesicle mediated transport. The association of apoptosis/cell death genes with hypertrophy would appear to further support a role for the increase in inflammatory genes identified herein. The identification of vesicle-mediated transport enriched genes is an intriguing potential mechanism for scWAT-mediated paracrine signaling to the myocardium. Future work will be needed to fully define whether increased inflammation or dysregulation of vesicular-mediated transport (or both) from scWAT is driving the phenotype seen in this model and to what extent these mechanisms play a broader role in the adipose-myocardial signaling axis.

The potential translational impact of decreased HuR expression in adipose tissue and its impact on CVD remain to be fully elucidated. Indeed, we observe a decrease in HuR expression in scWAT from obese wild-type mice, confirming the previously published report of decreased HuR expression in scWAT from obese humans (48). Here, we show that loss of HuR expression in adipose tissue is sufficient to induce CVD in the absence of obesity, but a direct mechanistic role for decreased adipose tissue expression of HuR in promoting CVD in humans remains unexplored. Future work will be needed to address this and other remaining questions. For example, HuR has been previously shown to be a mediator of inflammatory signaling (27, 42, 44, 71, 87), but the specific mechanisms by which loss of HuR expression in scWAT drives inflammatory signaling remains unknown. Furthermore, are scWAT-mediated increases in inflammatory mediators, such as TNF-αand IL-6 as we have shown here, sufficient to induce the observed CVD and to what extent might scWAT-derived extracellular vesicles signal to the myocardium? Another important remaining question is whether the loss of BAT-mediated thermogenesis directly contributes to the CVD observed in our model. An inverse clinical correlation between CVD and functional declines in BAT associated with aging, obesity, and other CVD co-morbidities has been noted, but the loss of functional BAT has yet to be conclusively shown to play a direct causal role in CVD.

## GRANTS

This work was partially supported by an American Heart Association Transformational Project Award (19TPA34910086; MT) and NIH grant R01 HL132111 (MT). LCG and SS are supported by AHA Predoctoral Fellowships (20PRE35210795 and 20PRE35230020, respectively).

## AUTHOR CONTRIBUTIONS

A.R.G., S.R.A., A.G., and M.T. conceived and designed the research. A.R.G., S.R.A., A.G., L.C.G., S.S., M.L.N., P.A., and M.T. performed experiments and analyzed data. A.R.G., S.R.A., A.G., J.B.B., O.K., and M.T. interpreted results. A.R.G., S.R.A., and M.T. drafted, edited, and revised the manuscript.

## FIGURE LEGENDS

